# Molecular basis of Ad5-nCoV Vaccine-Induced Immunogenicity

**DOI:** 10.1101/2024.06.21.600005

**Authors:** Dongyang Dong, Yutong Song, Shipo Wu, Busen Wang, Cheng Peng, Weizheng Kong, Zheyuan Zhang, Li-Hua Hou, Sai Li

## Abstract

In response to coronavirus disease 2019 (COVID-19), numerous vaccines have been developed to protect against SARS-CoV-2 infection. Ad5-nCoV (Convidecia) is a vaccine listed for emergency use by the WHO and has been administrated to millions of people globally. It comprises a series of human adenovirus 5 (Ad5) replication-incompetent vectored vaccines that transduce the spike protein (S) gene of various SARS-CoV-2 strains. Despite promising clinical data demonstrating its safety and effectiveness, the underlying molecular mechanism of its high immunogenicity and incidence of adverse reactions remains less understood. Here we combined cryo-ET, fluorescence microscopy and mass spectrometry to characterize the *in situ* structures, density and site-specific glycan compositions of the Ad5-nCoV_Wu and Ad5-nCoV_O vaccine-induced S antigens, which encode the unmodified SARS-CoV-2 Wuhan-Hu-1 S gene and optimized Omicron S gene, respectively. We found that the vaccine-induced S are structurally intact, antigenic and densely distributed on the cell membrane. Compared to Ad5-nCoV_Wu induced S, the Ad5-nCoV_O induced S demonstrate significantly better stability and is less likely to induce syncytia among inoculated cells. Our work demonstrated that Ad5-nCoV is a prominent platform for antigen induction and cryo-ET can be a useful technique for vaccine characterization and development.

## Introduction

SARS-CoV-2, the causative pathogen of COVID-19, is a single-stranded (+) RNA virus belonging to the β-coronavirus genus. The virus features rapid mutation rate and high infectivity. Among SARS-CoV-2 variants of concern (VOCs), the Omicron strain is still widely circulating in humans and mutating. With respect to Variants of Interest (VOI), the EG.5 strain is the most globally prevalent one to date, and XBB.1.5, XBB.1.16, BA.2.86, JN.1^1^ strains are also bothering a significant population. The SARS-CoV-2 spike (S) protein plays a critical role in viral infection and is the most important target for vaccine and antibody development. It comprises a S1 subunit, which contains a receptor binding domain (RBD) and is responsible for receptor binding with angiotensin-converting enzyme 2 (ACE2), and a S2 subunit functioning as a class-I fusion protein. During the maturation of S in ER/Golgi apparatus, its arginine-rich motif (PRRA) is cleaved by furin, a host enzyme protein, into S1 and S2 subunits^2^. An additional cleavage site S2’ is located near the fusion peptide. Upon S1 binding with ACE2, the exposed S2’ is cleaved by either transmembrane serine protease 2 (TMPRSS2) on the cell membrane or cathepsin L in the endosome for fusion activation^3^. In the fusion process, S undergoes extensive structural rearrangements, changing from a prefusion to a postfusion conformation^4^. The latter conformation contains only S2 and is non-immunogenic.

Up to now, the World Health Organization (WHO) has approved fourteen COVID-19 vaccines for emergency use^5^. These vaccines have been administrated globally and provided effective protection against severe illness, hospitalization and death from COVID-19. Technically, these vaccines are based on various platforms: two are based on mRNA, three based on inactivated virions, five based on protein subunits, and four based on adenoviral vectors. All fourteen vaccines chose S or its RBD as immunogens to stimulate adaptive immunity and help storing long-term immune memory. Ideally, immunogens brought out by vaccines are structurally intact and stable, abundant, and capable of stimulating sufficient immune responses. However, these requirements have been challenging particularly for SARS-CoV-2 vaccines. Firstly, compared to other viral class-I fusion proteins, SARS-CoV-2 S is structurally unstable. Such vulnerability results in the transformation of S into its non-immunogenic postfusion conformation and is especially problematic for the inactivated SARS-CoV-2 vaccines production^6^. Secondly, SARS-CoV-2 variants have been rapidly mutating and escaping the established protection offered by the previous vaccines. Genetic vaccines offer an efficient platform to overcome these difficulties by delivering the S genes into human cells and expressing the immunogens directly on the cell membrane. However, the cell-displayed S may trigger cell-cell fusion among neighboring cells and induce syncytia formation, which may potentially contribute to inflammatory response^7^, lung epithelium damage^8^, and immune dysfunction^8,9^. To mitigate these problems, 2P or 6P mutations and deletion of the S1/S2 cleavage site (PRRA) have been implemented into the design of licensed vaccines. For example, BNT162b2^10^, Ad26.COV2.S^11^and mRNA-1273^12^adopt the 2P mutation, NVX-CoV2373^13^ adopts both furin-cleavage site deletion and the 2P mutation, and vaccine candidates such as ChAdOx1 nCoV-19E6 adopt the 6P mutation^14^. Taken together, genetic vaccine platforms that enable prompt antigen sequence update and optimization may serve as an effective strategy to provide consistent protection against SARS-CoV-2 infection.

Adenoviral-vectored vaccines have been extensively developed against emerging viruses, such as Ad26-MVA against Ebola virus^15^, as well as Ad5-nCoV^16^, ChAdOx1 nCoV-19^17^ and Ad26.COV2-S^11^ against SARS-CoV-2. These vaccines utilize replication-defective adenovirus to transduce genes of immunogens into cells. Through expressing and displaying the immunogens on the membrane, these cells elicit humoral and cellular immunity^18^. Ad5-nCoV (Convidecia, CanSino Biologics) is a human adenovirus vector-based vaccine^19^ and is listed for emergency use by the WHO. The vaccine uses E1/E3 deleted Ad5 vectors to transduce wild-type SARS-CoV-2 S gene^20^. In an efficacy analysis conducted on 28 days post-vaccination adults, the Ad5-nCoV one-dose vaccine has demonstrated a 57.5% prophylactic efficacy against symptomatic COVID-19 and 91.7% against severe COVID-19 disease^21^. Building upon the prototype, the vaccine has been subsequently developed to encode SARS-CoV-2 Omicron S gene^22^. Ad5-nCoV had been approved in over ten countries and more than 100 million of doses were supplied worldwide during the COVID-19 pandemic. The aerosolized Ad5-nCoV was also been approved for emergency use in China in late 2022, which showed good immunogenicity as the booster vaccine and demonstrated effectiveness against infection in the real world^23,24^.

Despite the promising clinical data, the molecular basis of Ad5-nCoV induced potent immunogenicity is missing. A recent study on ChAdOx1 nCoV-19, a vaccine based on replication-deficient chimpanzee adenovirus vector, has reported the vaccine-induced S structures of WT SARS-CoV-2 and the Beta variant, along with corresponding glycan composition. These structures and glycan modifications mimic those found on the virions. Compared to the vaccine transducing WT S, the vaccine of HexaPro-stabilized (6P) S yields higher antigen expression, enhanced RBD exposure, and reduced S1 shedding^14,25^. In this work, we set out to characterize the structures and glycan compositions of two Ad5-nCoV vaccines encoding WT S or stabilized Omicron S, and compare them to those of ChAdOx1 nCoV-19. We determined the *in situ* structure, conformational variations and distribution of S by cryo-ET, and found that the syncytia formation can be mediated by Ad5-nCoV_Wu vaccine-induced S by fluorescence microscopy. We also performed site-specific N-linked glycan analysis of the vaccine-induced S by mass spectrometry and compared them to those of the native S on SARS-CoV-2 WT virions. Our work provides valuable information for the design and optimization of Ad5-nCoV. By examining the antigen expression and structural integrity, we also demonstrated the combination of fluorescent microscopy, cryo-ET and mass spectrometry provides molecular perspectives for the development of genetic vaccines against COVID-19 and other infectious diseases.

## Results

### Ad5-nCoV vaccines induce functional S protein on Vero cell membrane

Ad5-nCoV_Wu vaccine, which encodes the S gene of SARS-CoV-2 Wuhan-Hu-1 (WT) strain (NC_045512.2), and Ad5-nCoV_O vaccine, which encodes the S gene of SARS-CoV-2 Omicron strain B.1.1.529 (EPI_ISL_6640917) with 2P (K985P and V986P) mutation and S1/S2 cleavage site (PRRA) deletion (R682/R683/R685 deleted) (Fig. 1a), were used in this study. The Ad5-nCoV particles were produced and released from human embryonic kidney cells (HEK293), reaching a final concentration of 5×10^10^ viral particles per dose after purification and formulation. Negative staining electron microscopy (EM) examination verified that the vaccine particles are of high intactness and purity (Supplementary information, Fig. S1).

**Fig. 1.**
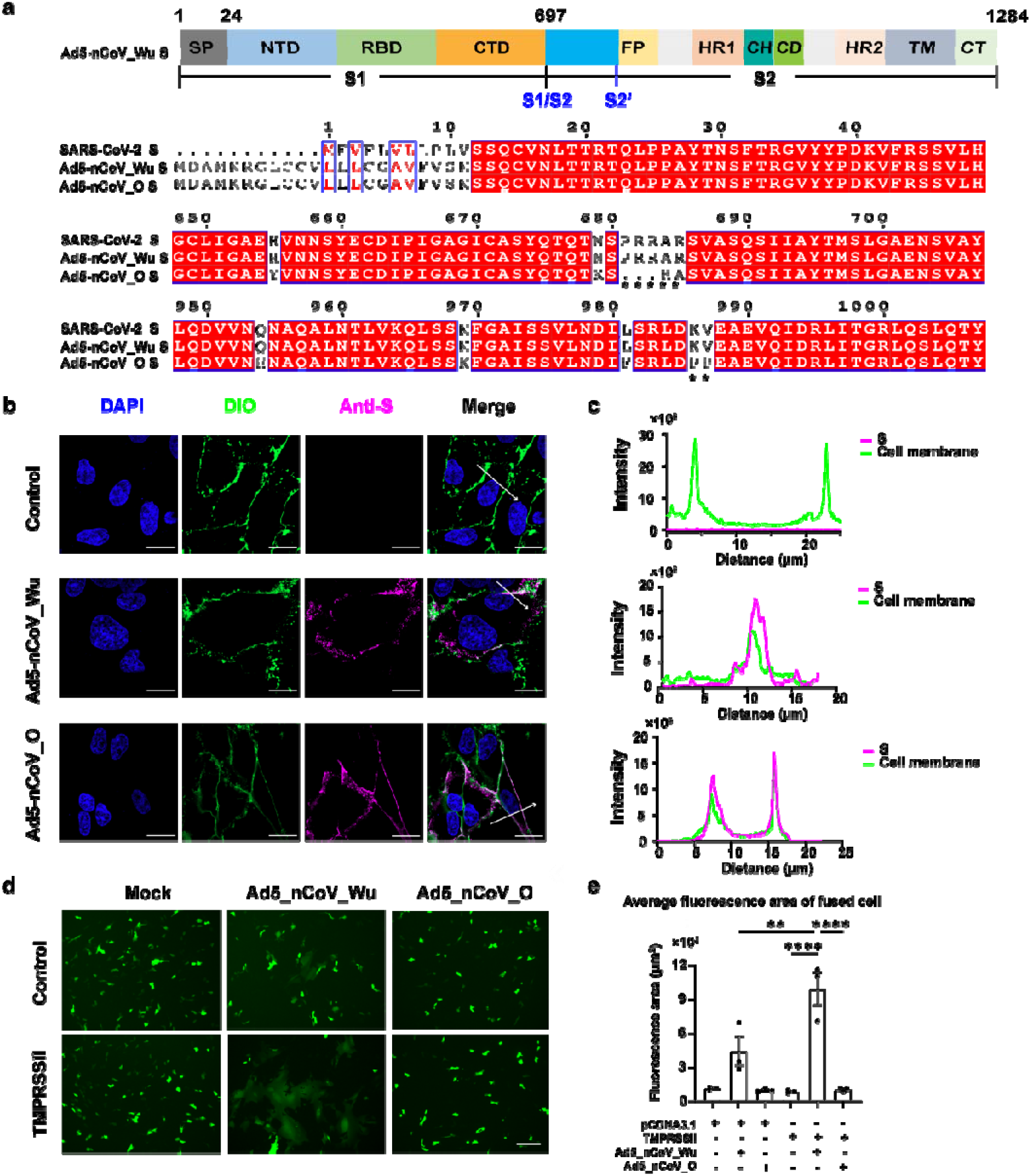
Light microscopy of Vero cells inoculated with Ad5-nCoV vaccines. **a** A schematic representation of S induced by the Ad5-nCoV_Wu vaccine. Compared to SARS-CoV-2 S, the signal peptide (SP) of Ad5-nCoV_Wu S is longer, but other domains remain the same. Abbreviations: NTD, N-terminal domain; RBD, receptor binding domain; CTD, C-terminal domain; S1/S2, S1/S2 cleavage site; S2’, S2’ cleavage site; FP, fusion peptide; HR1, heptad repeat 1; CH, central helix; CD, connector domain; HR2, heptad repeat 2; TM, transmembrane anchor; CT, cytoplasmic tail. The sequence alignment of S protein of SARS-CoV-2 with those of Ad5-nCoV vaccines is shown below. Two asterisk markers indicate the furin cleavage site deletion and 2P mutation on Ad5-nCoV_O S, respectively. **b** Immunofluorescent microscopy of Vero cells inoculated with two Ad5-nCoV vaccines. Cells were inoculated with Ad5-nCoV_Wu or Ad5-nCoV_O at MOI=1 for 48 h. The nuclei, membrane and vaccine-induced S were stained with DAPI (blue), DIO (green) and an anti-SARS-CoV-2-S1 antibody (magenta), respectively. Scale bar: 20 μm. **c** The fluorescence signal intensities of S (magenta) and membrane (green) along the white arrows in (**b**) were measured. **d** Fluorescence microscopy of vaccine-induced cell-cell fusion. Vero cells transfected with eYFP plasmids were inoculated with Ad5-nCoV vaccines, and then transfected with pcDNA3.1 or TMPRSS2 plasmids. The fluorescence images were captured at a 10× magnification. Scale bar: 200 μm. **e** The averaged fluorescence areas of fused cells in (**d**) were measured and analyzed with ImageJ (3 biologically independent samples per group). The P values were calculated using one-way ANOVA by GraphPad Prism 9 (****p<0.0001, **p<0.01).

To examine the expression level of S by Ad5-nCoV vaccines on the cell membrane, we first evaluated the subcellular localization of S induced by both vaccines using confocal microscopy. The Vero cell membrane was stained with a fluorescent membrane dye DIO (green) and S was stained with an antibody conjugated with a fluorescent label DyLight® 650 (magenta). As a result, S was detected on the cell membrane after Ad5-nCoV inoculations, but was not detected on Mock cells. The overlap of fluorescence intensity peaks along profiles spanning the cells evaluate the distribution of S on the cell membrane, revealing that both vaccines have induced an abundant amount of S (Fig. 1b, c).

Next, we evaluated the fusogenicity of Ad5-nCoV-induced S in mediating cell-cell fusion in the vaccine-inoculated Vero cells. We performed cell-cell fusion fluorescence assays in vitro and observed syncytia formation in the Ad5-nCoV_Wu-inoculated cells. In comparison, syncytia were not obvious in the Ad5-nCoV_O-inoculated cells or in the Mock cells (Fig. 1d, e). To test if the levels of TMPRSS2 present on the Vero cell membranes have affected the syncytia formation, TMPRSS2 was supplemented to the vaccine-inoculated cells by transfection, resulting in a nearly one-fold increase in the average area of syncytium in the Ad5-nCoV_Wu-inoculated cells. However, TMPRSS2 supplementation led to no obvious change in the levels of syncytia formation in the Ad5-nCoV_O-inoculated cells or in the Mock cells (Fig. 1d, e). To investigate if the syncytia formation is associated with S cleavage, we measured the levels of S in the vaccine-inoculated cells by Western blotting. S was detected in both Ad5-nCoV_Wu and Ad5-nCoV_O-inoculated cells regardless of TMPRSS2 supplementation, while S1 was undetectable in Ad5-nCoV_O-inoculated cells (Supplementary information, Fig. S2).

With the above evidence, we conclude that Ad5-nCoV vaccines are capable of inducing S on the cell membranes. The Ad5-nCoV_Wu induced S can be cleaved at the S1/S2 site. With S1/S2 cleavage, syncytia were abundantly observed in cell culture, suggesting that the Ad5-nCoV_Wu induced S is capable of mediating cell-cell fusion. In comparison, S1 shedding or syncytia were not observed in Ad5-nCoV_O-inoculated cells, suggesting that the 2P mutation and the S1/S2 cleavage site deletion are effective in stabilizing S in the prefusion conformation and thus preventing cell-cell fusion.

### Ad5-nCoV vaccines induce dense S on the cell membrane and extracellular vesicles

Next, we analyzed the *in situ* structures and distribution of S induced by the two Ad5-nCoV vaccines on cell membrane. Vero cells were seeded on EM grids, inoculated with the vaccines, plunge-frozen and subsequently imaged by cryo-ET. Examination of the cells revealed that the cell periphery was relatively thin, providing sufficient contrast for imaging and structural determination (Fig. 2a). Through scrutinizing the cell periphery, we observed densities of actin filaments, microtubules, intracellular vesicles and significant amounts of S-like particles (Fig. 2b, c). We also captured vesicles coated with S-like densities budding from (Fig. 2d), or in proximity to the cell periphery (Fig. 2b). To better illustrate the three-dimensional cell periphery, we reconstructed a composite structure of a representative filopodia by segmenting the densities of membrane, actin filaments and an S-coated vesicle, and projecting the S structures (see next session) onto their refined coordinates (Fig. 2e). With the above observations, we conclude that Ad5-nCoVs are capable of inducing high-density S on the inoculated cell membranes, and some of these S may subsequently relocate to cell-secreted extracellular vesicles (EVs).

**Fig. 2.**
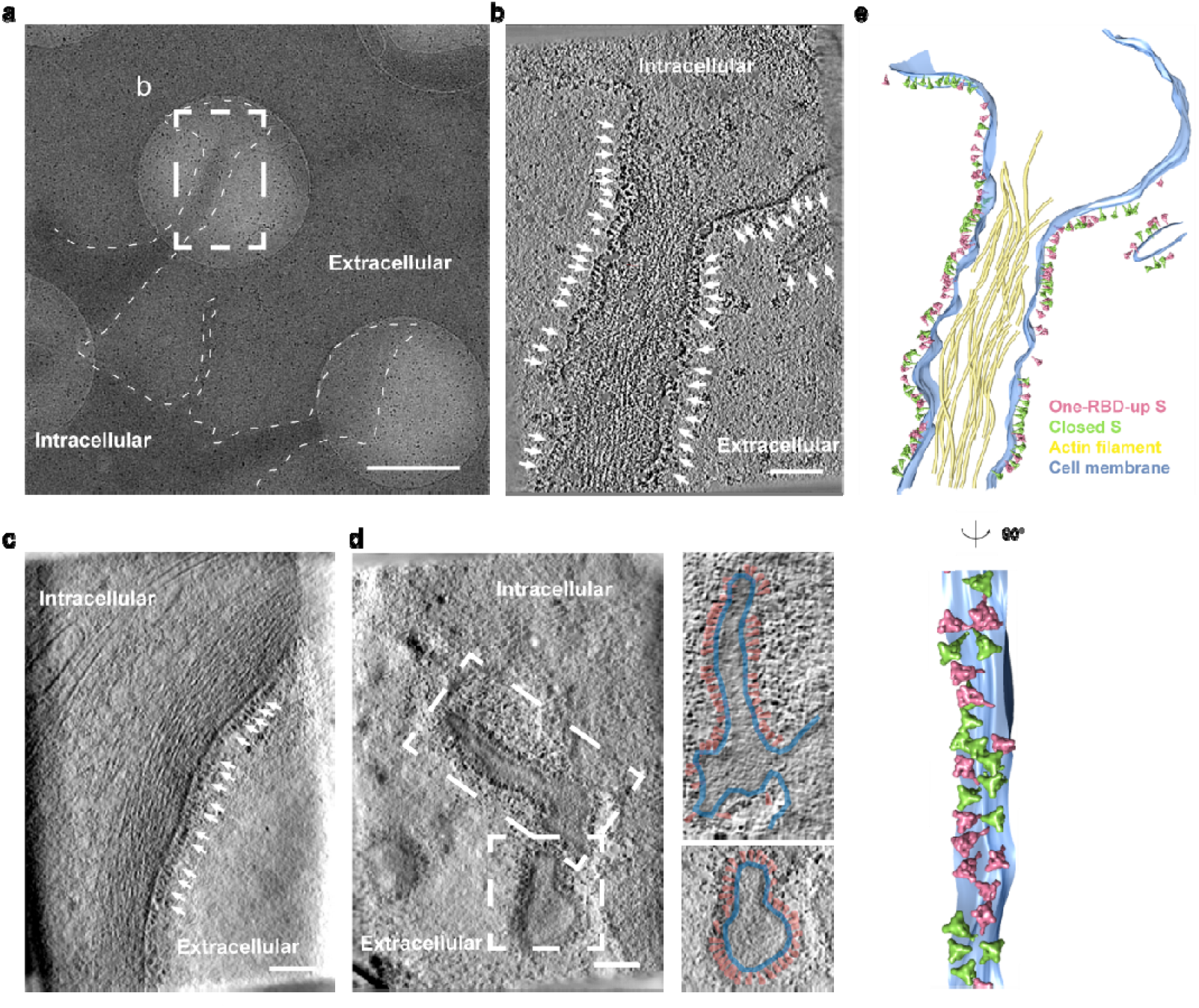
Cryo-ET of Vero cells inoculated with Ad5-nCoV vaccines. **a** A tomogram slice shows the filopodia of a cell inoculated with Ad5-nCoV_Wu vaccine. The thin dashed line outlines the cell periphery. Scale bar: 1 μm. **b** A zoomed-in view of the dashed box in (**a**). White arrows indicate densities corresponding to the Ad5-nCoV_Wu-induced S. Tomogram thickness: 7.8 nm. Scale bar: 100 nm. **c** The top- and side view of the filopodial structure in (**b**). The structure is reconstructed by segmenting the membrane (blue) and actin filament (yellow) from the corresponding densities and projecting the closed (green) and one-RBD-up (pink) S structures back onto their refined coordinates. **d** A tomogram slice shows S densities (white arrows) on the periphery of an Ad5-nCoV_O-inoculated cell. Tomogram thickness: 16.4 nm. Scale bar: 100 nm. **e** The left panel is a tomogram slice showing two extracellular vesicles (dashed boxes) in the vicinity of an Ad5-nCoV_O-inoculated cell. The right panels are display the magnified views, corresponding to the dashed boxes in the left panel, with specific highlights for S (pink) and the vesicle membrane (blue). Tomogram thickness: 3.12 nm. Scale bar: 100 nm.

### The Ad5-nCoV vaccine-induced S predominantly adopt prefusion conformation, exhibit antigenicity and interact with antibodies

To determine the identity of the dense S-like particles present on the cell membrane, we annotated 1,739 and 2,281 S-like particles on the membrane of Ad5-nCoV_Wu and Ad5-nCoV_O-inoculated cells, respectively (Supplementary information, Table S1). Subtomogram averaging of these particles has revealed that they were structurally intact, predominantly in prefusion conformation (Fig. 3a), and fit well with the structure of a predicted full-length S^26^ (Supplementary information, Fig. S3). Subsequent classification revealed that 51.5% of prefusion S adopted the closed conformation and 48.5% adopted the one-RBD-up conformation on the Ad5-nCoV_Wu-inoculated cells. In comparison, 22% of prefusion S adopted closed conformation and 78% of S adopted one-RBD-up conformation on the Ad5-nCoV_O-inoculated cells (Fig. 3b). The conformational distributions of S were similar to that observed on SARS-CoV-2 WT virions^27^ and that of the recombinantly expressed full-length Omicron S^28^, respectively. Notably, we did not distinguish postfusion S from the cell membrane.

**Fig. 3.**
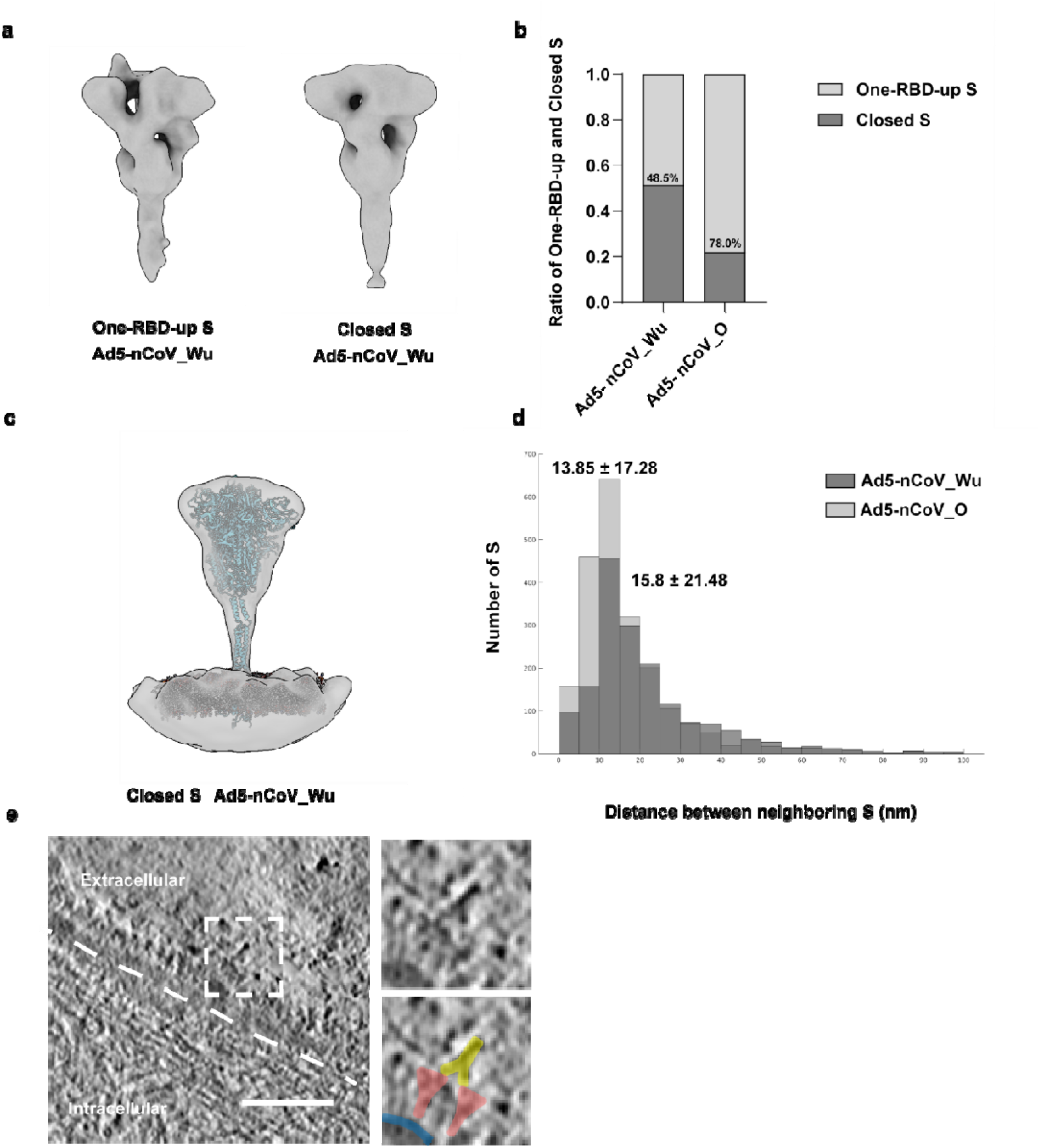
The native structure and the distribution of Ad5-nCoV induced S. **a** The native structure of the Ad5-nCoV_Wu-induced S in one-RBD-up or closed conformations. **b** The ratio of the one-RBD-up and the closed S induced by Ad5-nCoV_Wu and Ad5-nCoV_O. **c** The closed Ad5-nCoV_Wu S map shows the density of its stalk region and the adjoining membrane. A full-length model of S embedded in a lipid bilayer^26^ is fitted to the map for comparison. **d** A histogram shows the distance between the closest S proteins induced by Ad5-nCoV_Wu or Ad5-nCoV_O. **e** The left panel is a tomogram slice showing CV3-13 IgG molecules bound to Ad5-nCoV_O-induced S proteins. The right panels display the magnified views, corresponding to the dashed box in the left panel, with specific highlights for S (red), IgG (yellow), and the cell membrane (blue). Tomogram thickness: 1.64 nm. Scale bar: 100 nm.

With the refined structures and coordinates, we next analyzed the density of S on the two vaccine-inoculated cells. The stalk regions of the closed-conformation S induced by Ad5-nCoV_Wu and Ad5-nCoV_O were both 9.4 nm in length, 2.5 nm longer than that on the SARS-CoV-2 WT^27^ virions (Supplementary information, Fig. S3). By fitting a predicted full-length model of closed S^26^, we confirmed that the density at the lower end of the Ad5-nCoV-induced S indeed corresponded to an erected, full-length stalk (Fig. 3c). We suspect that the perpendicular feature of the Ad5-nCoV-induced S stalk region, which is absent from the native S structures determined on SARS-CoV-2, is attributed to the crowding of S on the cell membrane that restricts the flexibility of the S hinge. We further measured the distance between the nearest S particles, revealing an average distance of 15.8 nm for Ad5-nCoV_Wu S and 13.85 nm for Ad5-nCoV_O S (Fig. 3d). To investigate the antigenicity of S on the cell membrane, we tested whether they interact with non-neutralizing antibodies. The vaccine-inoculated cells were incubated with CV3-13 Ab, an antibody with potent Fc-mediated effector functions^29^, and then imaged by cryo-ET. Densities corresponding to S-IgG complexes were discernable on the cell membrane (Fig. 3e).

Together, these observations suggest that the Ad5-nCoV-induced S are structurally intact, predominantly in prefusion conformation, and densely distributed on the cell membrane. Compared to the Ad5-nCoV_Wu S, the Ad5-nCoV_O S were denser, with a higher proportion adopting the one-RBD-up conformation. Moreover, the induced S proteins can be recognized by SARS-CoV-2 specific Abs, suggesting that these S are antigenic.

### Site-specific glycan analysis of the Ad5-nCoV vaccine-induced S

Glycan modifications on S facilitate protein folding. By shielding certain epitopes, glycans also aid the virus in immune-evasion. Therefore, glycosylation of the vaccine-induced antigens plays important role in determining the efficiency of the vaccine-induced antibodies in targeting the exposed epitopes of authentic viral antigens and thereby providing prophylaxis. To analyze the site-specific glycan modifications on Ad5-nCoV-induced S on the cellular membrane, we purified the S proteins from vaccine-inoculated HEK293F cells by pull-down assays (Supplementary information, Fig. S4). The bands corresponding to S proteins were cut out from SDS-PAGE gel, digested with protease and analyzed by liquid chromatography-mass spectrometry (LC/MS).

Based on branching and fucosylation, the glycans were classified into four types: core, oligo-mannose, hybrid and complex. Overall, the glycosylation on Ad5-nCoV expressed S were consistent with those of the native SARS-CoV-2 WT S^27^(Fig. 4). More complex-typed glycans were identified on Ad5-nCoV_O S (66%, Supplementary information, Table S2) than on Ad5-nCoV_Wu S (Fig. 4). We analyzed the vaccines-expressed S based on their sequences starting from the signal peptide. Different variants of S have different protein sequences and signal peptides. Therefore, the sequence number of each glycan site on the vaccines is slightly different from that of the WT viral S. Notably, the N242 of Ad5-nCoV_Wu S and N245 of Ad5-nCoV_O S showed high degrees of oligo-mannose-type glycosylation, which is consistent with N234 of the native SARS-CoV-2 WT S. Glycosylation at this specific site has been reported to modulate the conformational transition of RBD^26,30,31^. The ratio of complex-typed glycans on Ad5-nCoV_Wu S (56%, Supplementary information, Table S2) was slightly lower than that of the native SARS-CoV-2 WT S. We have detected more glycosylation (19 sites of Ad5-nCoV_Wu S and 23 of Ad5-nCoV_O S) and higher proportion of complex-typed glycans on Ad5-nCoV-induced S (55.5% of Ad5-nCoV_Wu S and 66% of Ad5-nCoV_O S, Supplementary information, Table S2) than that of the ChAdOx1 nCoV-19 expressed S (18 glycosylation sites; 15% are complex)^25^. Taken together, these data suggest that the glycans on Ad5-nCoV-induced S are more mature, providing a more similar glycan profile to that of native viral S.

**Fig. 4.**
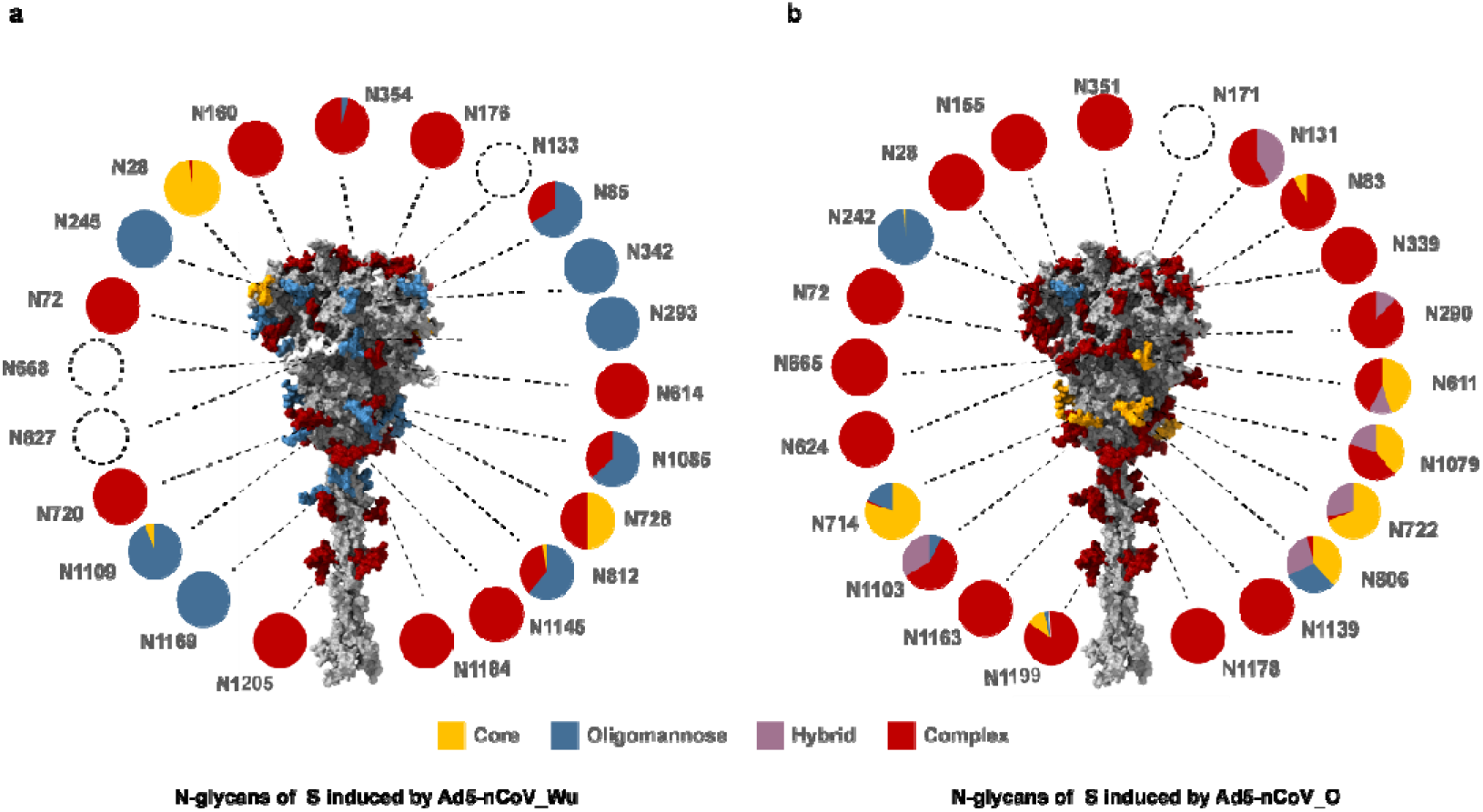
Site-specific glycan composition of the vaccine-induced S. The N-linked glycan composition of S proteins, induced by Ad5-nCoV_Wu (**a**) and Ad5-nCoV_O (**b**), were analyzed by MS and are represented in pie charts. The dashed circles represent the undetected glycans. The 22 glycans are highlighted on a full-length model^26^, with each glycan being colored according to their compositions. In case of mixed compositions, the glycan is colored according to the component that constitutes the highest percentage. The glycans on vaccines-induced S were analyzed based on their sequences starting from the signal peptide. Due to the different protein sequences and signal peptides from the WT viral S, the sequence number of each glycan site on the vaccines-induced S has slight offset compared to that of the WT viral S.

## Discussion

Structural integrity, abundance, RBD-rising dynamics are key factors influencing the immunogenicity of vaccine-induced antigens. However, various ratios of postfusion S have been observed on infectious SARS-CoV-2 virions^32^, formaldehyde-inactivated intact SARS-CoV-2 WT^27^, D614G^33,34^ and Delta^35^ virions, electron beam-irradiated Delta^35^ virions and β-propiolactone (BPL)-inactivated WT virions (Supplementary information, Table S3), with the BPL-inactivated virions harboring the highest ratio of postfusion S (74.4%)^36^. These observations indicate that the SARS-CoV-2 S is susceptible to both chemical and physical stresses. This poses challenges for the production of inactivated SARS-CoV-2 vaccines, which predominantly utilizes BPL as the inactivating agent^37^. Apart from structural integrity, antigen abundance is another factor in eliciting immune response. Most SARS-CoV-2 specific antibodies target RBD, of the four major classes of SARS-CoV-2 RBD-nAbs, class-1 and class-4 nAbs only bind to up-RBD, while class-2 and class-3 nAbs bind both up- and down-RBDs^38^. RBDs that possess rising dynamics expose more conserved epitopes and increase the production of cross-reactive neutralizing antibodies against a variety of strains^39^. Therefore, more proportions of raised RBD and dynamics of RBD rising increases the immunogenicity of S and is beneficial for vaccine effectiveness. To evaluate these molecular aspects of Ad5-nCoV-induced antigens, we determined the *in situ* structures and glycan compositions of vaccine-induced S on the cell membrane. Our results show that their structures, conformational ratio and glycosylation mimic those of the authentic virions. Cryo-ET has further confirmed the binding between S on Ad5-nCoV_O-inoculated cells and CV3-13 Ab, suggesting that these vaccine-induced S are antigenic. With respect to antigen abundance, we showed that S proteins induced by Ad5-nCoV_Wu and Ad5-nCoV_O vaccines display densely on the cell membrane and on the EVs. It has been shown that the antigen-coated EVs budded from antigen-presenting cells (APC) can present antigen peptides to and activate T cells. In return, EVs can also transfer antigen peptides to APC to prime naive T or B cells for activation^40^. Therefore, the metastatic property of these S-coated EVs broadens the distribution of vaccine-induced antigens^41^. Finally, we showed that the 2P mutation and furin cleavage site deletion are effective in stabilizing the prefusion conformation for enhanced immunogenicity of Ad5-nCoV_O.

Despite the good safety profile exhibited on the aerosolized Ad5-nCoV, its recipients had slightly higher incidence of adverse reactions than those who received inactivated COVID-19 vaccine^24^. To explore the possible mechanism of the adverse effects, we applied immunofluorescence microscopy (IFM) experiments to investigate the syncytia formation among inoculated cells. Upon cleavage at the S1/S2 site, the interaction between S and ACE2 would trigger S1 shedding and induce the cell-cell fusion between inoculated cells and their neighboring cells^42^. This is obvious in Ad5-nCoV_Wu-inoculated cells, in which syncytia were observed and cleaved S1 subunits were detected in the supernatant. In contrast, cells inoculated with Ad5-nCoV_O, which encodes S with 2P mutation and furin-cleavage site deletion, did not exhibit obvious syncytialization and S1 shedding. Also, our study demonstrated that the syncytia formation can be facilitated by TMPRSS2. Both ACE2 and TMPRSS2 have been reported to present on the nasal epithelium, airway epithelium, lung epithelium and esophagus^43–45^, and SARS-CoV-2 S-induced pneumocytes fusion has been reported to cause nuclear damage, micronuclei formation, a type I interferon (IFN) response enhancement and cytokine production, which could exacerbate illness^46^. These effects can be alleviated by administering Ad5-nCoV_O vaccine, while Ad5-nCoV_Wu would not cause severe syncytia if administered properly. Given ACE2 and TMPRSS2 are rarely present on muscle tissue^43–45^, and the Ad5-nCoV vaccine can be administered by either intramuscular injection^19^ or inhalation^47,48^, the Ad5-nCoV_Wu shall be administered by intramuscular injection to avoid syncytialization.

Altogether, our results indicate the antigenicity of the Ad5-nCoV-induced S was robustly corroborated, not only by the authentic prefusion structure, the dynamics of RBD rising and glycosylation, but also by their accessibility and distribution. These observations potentially explain for the significantly higher and longer lasting levels of induced neutralizing antibodies against SARS-CoV-2 and better protection against SARS-CoV-2 infection by Ad5-nCoV than those did by inactivated COVID-19 vaccine^24^. We also showed that cryo-ET can be a useful technique in vaccine evaluation and development. Building on these findings, the adenoviral vectored vaccine, recognized for its efficient packaging and potent immunogenicity and versatile delivery routes, demonstrates its potential in offering protection against highly mutable viruses that pose a threat to human health, including SARS-CoV-2 and potential future emerging viruses.

## Materials and Methods

### Ad5-nCoV vaccine production

The Ad5-nCoV vaccines were developed by Beijing Institute of Biotechnology and CanSinoBIO. Ad5-nCoV_Wu encoding the S protein of the Wuhan-Hu-1 strain (NC_045512.2) without any mutations^49^, while Ad5-nCoV_O encoding the S protein of the B.1.529 strain (EPI_ISL_6640917) with the furin cleavage site mutation (R682/R683/R685 deletions) and two proline substitutions at residues K986 and V987^50^. The gene of the S proteins were codon optimized, and the signal peptides (aa 1-13) were replaced by tPA. The vaccines were constructed with the AdMax adenovirus system (Microbix Biosystem, Canada), propagated in HEK 293 cells, and purified by ion-exchange and size-exclusion chromatography. The purified vaccines contained 5×10^10^ particles per dose.

### Negative staining electron microscopy

5 μL vaccine solution (1×_10^11^ vp/ml) was loaded onto the surface of the carbon-coated glow-discharged copper grid for 2 minutes. Subsequently, samples were stained with 3 μL of 1% uranyl acetate. Digital micrographs were then captured at 120 kV using Tecnai Spirit TEM (Thermo Fisher Scientific, Hillsboro, OR).

### Immunofluorescence microscopy

Vero cells (ATCC CCL-81) were cultured in 35 mm confocal dishes (D35C4-20-1-N) with DMEM (Gibco, Carlsbad, CA) supplemented with 5% FBS (Gibco, Carlsbad, CA) and 1% Pen/Strep (Gibco, Carlsbad, CA). When reached 60% confluency, the cells were inoculated with Ad5-nCoV_Wu or Ad5-nCoV_O vaccine at MOI = 1 for 48 h. Vaccine-inoculated cells and control cells were rinsed with phosphate-buffered saline (PBS, Gibco, Carlsbad, CA), and fixed with 4% paraformaldehyde (Sino Biological. Inc., Beijing, China). SARS-CoV-2 S proteins induced by vaccines on the cell membrane were recognized by S309 primary antibody. The primary antibodies were subsequently recognized by a goat anti-human IgG secondary antibody conjugated with DyLight® 650 fluorescent label (Abcam, Cambridge, UK). Cell nuclei were stained with DAPI (Beyotime. Inc., Shanghai, China). Cell membrane was stained with DIO (Beyotime. Inc., Shanghai, China). Samples images were then captured with Zeiss LSM980 Airyscan2 confocal microscope using a 100× oil immersion objective lens (ZEISS, Oberkochen, Germany). The images and the fluorescence intensity along profiles spanning the cells were analyzed using ZEN3.0 software.

### Fluorescence microscopy

Vero cells were cultured in 6-well plates as described previously. At 60% confluency, cells were transfected with plasmids encoding eYFP and inoculated with Ad5-nCoV_Wu (MOI = 1), Ad5-nCoV_O (MOI = 1), or left untreated. 48 h post-vaccination, all types of cells were transfected with pcDNA3.1 or plasmids encoding TMPRSS2. The eYFP fluorescence signals were examined and captured using the fluorescence microscope EVOS™ M5000 (Thermo Fisher Scientific, Hillsboro, OR) at a 10× magnification. The average area of cells or syncytium was measured using ImageJ software (3 biologically independent samples per group). The P values were calculated using one-way ANOVA by GraphPad Prism 9 (****p<0.0001, **p<0.01). After imaging, cells were collected for western blotting.

### Western blotting

Vero cells from the 6-well plates were resuspended in 200 μL RIPA buffer (Beyotime Biotec. Inc., Shanghai, China) containing 0.1 mM Phenylmethylsulfonyl fluoride (PMSF). The cells were then incubated on ice for at least 10_min to ensure complete cell lysis. Preliminary protein quantification was carried out using BCA protein assay (Cowin Biotech Co., Ltd, Beijing, China). Cell lysate was then loaded onto an SDS-PAGE gel (GenScript Biotech Corporation, Jiangsu, China) and transferred to a polyvinylidene fluoride (PVDF) membrane (Merck Millipore Ltd., Co. Cork, Ireland). Rabbit anti-S1 polyclonal antibody (Sino biological, Inc., Beijing, China) and mouse anti-β-actin antibody (Sino Biological, Inc., Beijing, China) were used as primary antibodies. ImageJ 1.45 software (National Institutes of Health, Bethesda, MD) was used for protein quantification.

### Cryo-ET sample preparation and imaging

For Ad5-nCoV_Wu sample, gold grids coated with holey carbon film (300 mesh, R2/2, Quantifoil, Jena, Germany) were glow-discharged and UV-treated for 1 h. And after 20 min treatment with 20 μg/mL bovine fibronectin (Merck millipore Ltd., Co. Cork, Ireland), the grids were washed with PBS. 1 × 10^5^ Vero cells were seeded onto the grids in 35 mm dishes and cultured in DMEM supplemented with 10% FBS and 1% Pen/Strep. The grids were incubated overnight at 37°C, 5% CO_2_ to enhance cell adhesion. The cells were then inoculated with Ad5-nCoV_Wu vaccine at MOI = 20 and incubated at 37 °C, 5% CO_2_ for 48 h. Afterwards, grids were applied with 3 μL 10 nm diameter BSA Gold Tracer (Aurion, The Netherlands) and single-side blotted using the EMGP plunger (Leica, Wezlar, Germany). Subsequent imaging of vitrified grids was performed on a Titan Krios microscope (Thermo Fisher Scientific, Hillsboro, OR) operated at 300 kV equipped with a K3 direct electron detector (Gatan Inc., CA). 28 tilt series were acquired in super-resolution mode at a nominal magnification of 18,000 ×, resulting in a calibrated pixel size of 0.78 Å. Data were collected using the dose-symmetric scheme from −60° to 60° at 3° intervals with a defocus range from −4 to −5 μm in SerialEM^51^. 8 frames were recorded per tilt and the total dose of each tilt series was 131.2 e^−^/Å^2^.

For Ad5-nCoV_O sample, Vero cells were seeded either on gold finder grids (200 mesh, R2/2, Quantifoil, Jena, Germany) or gold grids coated with lacey carbon film (XXBR Technology, Beijing, China) and infected with Ad5-nCoV_O vaccine. The culture, infection and vitrification procedures were consistent with Ad5-nCoV_Wu sample. Vitrified grids were imaged on a Titan Krios microscope (Thermo Fisher Scientific, Hillsboro, OR) operated at 300 kV equipped with a Gatan BioQuantum energy filter (slit width 20 eV, Gatan, CA) and K3 direct electron detector (Gatan, CA). 70 tilt series were acquired in super-resolution mode at a nominal magnification of 53,000 ×, resulting in a calibrated pixel size of 0.83 Å. Data were collected using the dose-symmetric scheme from −60° to 60° at 3° steps with a defocus range from −3.4 to −5.6 μm in SerialEM^51^. At each tilt, 8 frames were recorded and the total dose of each tilt series was 106.6 e^−^/Å^2^.

For Ad5-nCoV_O + CV13_3 sample, Vero cells were seeded on grids and were inoculated with Ad5-nCoV_O. At 48 h post infection, anti-RBD antibody CV13_3 was added at a final concentration of 50 μg/mL and incubated for 6 h. Other sample preparation and imaging details were same as those of Ad5-nCoV_O sample.

### Cryo-ET data processing

All tilt series data were processed in a high-throughput preprocessing suit developed within our lab^27^. Motion between frames at each tilt were corrected using MotionCor^52^ and MotionCor2^53^. Defocuses of tilt series were estimated using Gctf^54^. After tilt series alignment in IMOD^55^, 22 Ad5-nCoV_Wu tilt series and 27 Ad5-nCoV_O tilt series with good fiducial alignment and evident spike features were kept for following processing. Tomograms were contrast transfer function corrected and reconstructed by weighted back projection in NovaCTF^56^, resulting in final pixel sizes of 1.56 Å/pixel for Ad5-nCoV_Wu and 1.66 Å/pixel for Ad5-nCoV_O, respectively. For better visualization, 8 × binned tomograms were missing wedge corrected using IsoNet^57^. 1,739 Ad5-nCoV_Wu spikes and 2,281 Ad5-nCoV_O spikes were identified manually from the denoised 8 × binned tomograms.

Subtomogram averaging were performed in Dynamo^58^. For Ad5-nCoV_Wu sample, 1,739 subtomograms with a box size of 36 × 36 × 36 were extracted from 8 × binned tomograms and the cropped subtomograms were averaged to generate an initial template. Resolution was restricted to 35 Å and C3 symmetry was applied at this stage. The refined coordinates were used to reextracted subtomograms from 4 × binned tomograms with a box size of 72 × 72 × 72 for further alignment, where C3 symmetry was applied and resolution was restricted to 25Å. The aligned particles were subjected to a multi-reference classification using 35 Å low-pass filtered closed S and one-RBD-up S from WT SARS-CoV-2 virions (EMD-30426, EMD-30427)^27^ as templates. C1 symmetry was applied during alignment. After classification, 894 closed S and 843 one-RBD-up S were distinguished and their subtomograms were reextracted from 2 × binned tomograms with a box size of 144 × 144 × 144. Further alignments of 2 × binned S used a customized “gold-standard adaptive bandpass filter” method^27^, and the resolution was estimated using a 0.143 criterion for the Fourier shell correlation. C3 symmetry and C1 symmetry were applied for closed S and one-RBD-up S, respectively. Finally, a 14.0 Å map of closed S and a 22.5 Å map of the one-RBD-up S were achieved. To display the density connecting the spike and membrane, the refined coordinates and orientations of the closed S were transferred to RELION4.0 for further reconstruction, as shown in Fig. 3c.

For Ad5-nCoV_O sample, 2,281 subtomograms with a box size of 36 × 36 × 36 were extracted from 8 × binned tomograms. The aligned result of 8 × binned Ad5-nCoV_Wu S was used as template. Resolution was restricted to 45 Å and C3 symmetry was applied at this stage. The refined coordinates were used to reextracted subtomograms from 4 × binned tomograms with a box size of 72 × 72 × 72 for further alignment, where C3 symmetry was applied and resolution was restricted to 25Å. The aligned particles were subjected to a multi-reference classification using 35 Å low-pass filtered closed S and one-RBD-up S from SARS-CoV-2 virions (EMD-30426, EMD-30427)^27^ as templates. C1 symmetry was applied during this alignment. After classification, 501 closed S and 1,778 one-RBD-up S were distinguished and their coordinates were reextracted from 2 × binned tomograms with a box size of 144 × 144 × 144. Further alignments of 2 × binned S used a customized “gold-standard adaptive bandpass filter” method^27^, and the resolution was estimated using a 0.143 criterion for the Fourier shell correlation. C3 symmetry and C1 symmetry were applied for closed S and one-RBD-up S, respectively. Finally, an 18.4 Å map of closed S and a 17.1 Å map of the one-RBD-up S were achieved.

### Antibody expression and purification

The encoding regions of the variable heavy (VH) and variable light chains (VL) of antibodies were synthesized by Tsingke (Tsingke, Beijing, China). These two regions of antibodies were cloned into the AbVec2.0-IGHG1 and AbVec1.1-IGKC vector, respectively. IgG antibodies were transiently expressed in HEK 293F cells (1.8 × 10^6^ cells/mL) using 1 μg of total DNA per million cells, with a ratio of 1:2:9 for light chain plasmid, heavy chain plasmid, and PEI (Polysciences, Inc., Warrington, PA). After incubation for 72 h at 37 °C with 5% CO_2_ and 125 rpm oscillation, cell suspensions were harvested by centrifugation at 3000 rpm for 15 min. The supernatant was then filtered using a 0.45 μm filter (Millipore Ltd., Co. Cork, Ireland). Antibodies were purified by affinity chromatography using a HiTrap Protein A HP column (Cytiva, Logan, UT). Fractions containing the protein were concentrated and further purified by size-exclusion chromatography using a Superdex 200 increase 10/300 GL column (Cytiva, Logan, UT).

### Pull-down assay of full-length S proteins induced by two Ad5-nCoV vaccines

Ad5-nCoV vaccines were administered to HEK 293F cells at MOI=1. The cells were incubated for 48 h at 37 °C and 120 rpm oscillation. The cell pellets were then collected by centrifugation at 3000 rpm for 15 min. Cell pellets were resuspended in HEPES buffer (25 mM HEPES 7.4 and 150 mM NaCl) and subjected to ultrasonic lysis. The lysate was ultra-centrifugated at 40,000 rpm for 1 h, and the resulting pellets were treated with 2% DDM (Inalco Pharm, San Luis Obispo, CA) dissolved in HEPES buffer. The S proteins anchored at cell membrane, extracted by DDM, were ultra-centrifugated at 40,000 rpm for 1 h. The resulting supernatant containing the S proteins was incubated with S309 antibody in a sodium phosphate buffer containing 0.17% DDM. The S-antibody complexes were purified with protein-A affinity columns (Cytiva, Logan, UT).

### Mass spectrometric analysis

The S-S309 antigen-antibody antibody complexes were electrophoresed on the 4-12% SurePAGE™ Bis-Tris gel (Genscript Biotech Corporation, Jiangsu, China). The proteins were visualized by One-Step Blue Stain (Biotium, Fremont, CA). Bands of S protein were excised from the gel and processed as previously described^27^. In brief, the bands were reduced with 5 mM DTT, alkylated with 11 mM iodoacetamide and digested with Trypsin Gold (Promega, Madison, WI), Chymotrypsin (Promega, Madison, WI), Trypsin Gold and Alpha Lytic protease (Sigma-Aldrich, St. Louis, MO), or only alpha Lytic protease in 50 mM ammonium bicarbonate at 37 _ overnight. After digestion, samples were quenched with 10% trifluoroacetic acid (TFA) to adjust the pH below 2. Peptides were extracted with 0.1% TFA in 50% acetonitrile aqueous solution for 1 h and followed by vacuum drying in speedVac. The peptides above were redissolved in 25 μL 0.1% TFA and 6 μL of them were analyzed by Orbitrap Exploris 480 mass spectrometer (Thermo Fisher Scientific).

For LC-MS/MS analysis, the peptides were separated by a 60 min gradient elution at a flow rate of 0.30 µL/min with Thermo-Dionex Ultimate 3000 HPLC system, which was directly interfaced with an Orbitrap Exploris 480 mass spectrometer (Thermo Fisher Scientific, Hillsboro, OR). The analytical column was a homemade fused silica capillary column (75 µm ID, 150 mm length; Upchurch, Oak Harbor, WA) packed with C-18 resin (300 Å, 5 µm, Varian, Lexington, MA). Mobile phase A consisted of 0.1% formic acid, and mobile phase B consisted of 100% acetonitrile and 0.1% formic acid. The Orbitrap Exploris 480 mass spectrometer was operated in the data-dependent acquisition mode using Xcalibur 4.3.73.11 software and there was a single full-scan mass spectrum in the orbitrap (350-1,550 m/z, 120,000 resolution) followed by top-speed MS/MS scans in the Orbitrap.

Glycopeptide fragmentation data were extracted from the raw file using Byonic™ (Version 2.8.2). The data were searched with N-glycan 309 mammalian no sodium library. The search criteria were as follows: non-specificity, three missed cleavages were allowed; the oxidation (M) and 54.01063 Da (F) were set as the variable modification; precursor ion mass tolerances were set at 20 ppm for all MS acquired in an orbitrap mass analyzer; and the fragment ion mass tolerance was set at 0.02 Da for all MS2 spectra acquired. The peptide false discovery rate (FDR) was calculated using Fixed value PSM validator provided by PD. When the q value was smaller than 1%, the peptide spectrum match (PSM) was considered to be correct. FDR was determined based on PSMs when searched against the reverse, decoy database. Peptides only assigned to a given protein group were considered as unique. The false discovery rate (FDR) was also set to 0.01 for protein identifications.

As in the previous study^27^, the data with a score less than 30 were excluded. The N-glycoform abundance at each site was analyzed according to intensities. The glycans were classified into oligomannose type, hybrid type, complex type, and core type. Hybrid and complex glycans were further classified according to fucose components and antennas types. The ratio of each glycan type was determined by calculating the mean of the three replicates.

## Data and materials availability

Electron microscopy maps have been deposited in the Electron Microscopy Data Bank under accession codes EMD-XXXXX and EMD-XXXXX. The mass spectrometry proteomics data have been deposited to the ProteomeXchange Consortium via the PRIDE^59^ partner repository with the dataset identifier PXDXXXXX.

## Acknowledgements

We thank Dr. Jianlin Lei, Dr. Fan Yang and Dr. Xiaomin Li from the cryo-EM Facility, Technology Center for Protein Sciences, Tsinghua University, for their support on cryo-EM data collection. We thank the computational facility support on the cluster of Bio-Computing Platform (Tsinghua University Branch of China National Center for Protein Sciences Beijing). We thank Xiaolin Tian and Dr. Haiteng Deng in Technology Center for Protein Sciences, Tsinghua University, for MS analysis. We thank the assistance of Imaging Core Facility, Technology Center for Protein Sciences, Tsinghua University.

## Funding

This work was supported in part from National Natural Science Foundation of China #32241031 and #32171195, Tsinghua University Spring Breeze Fund #2021Z99CFZ004 and Dushi Fund #2023Z11DSZ001.

## Author Contributions

S.L. conceived and designed the project. S.W, B.W. and L.H. prepared the Ad5-nCoV sample. D.D. performed the fusion assay, immunofluorescence microscopy, purified the antibody and S-antibody complex and prepared the sample for cryo-ET. D.D. and Y.S. collected the cryo-ET data. Y.S., C.P., D.D. and Z.Z. analyzed the cryo-ET data. D.D. and Y.S. analyzed the glycan data. D.D., Y.S., W.K., L.H. and S.L. wrote the manuscript. All authors critically revised the manuscript.

## Conflict of Interest

The authors declare no competing interests.

## Notes

### Competing Interest Statement

The authors have declared no competing interest.

