## Supplementary Info for "Molecular basis of Ad5-nCoV Vaccine-Induced Immunogenicity"

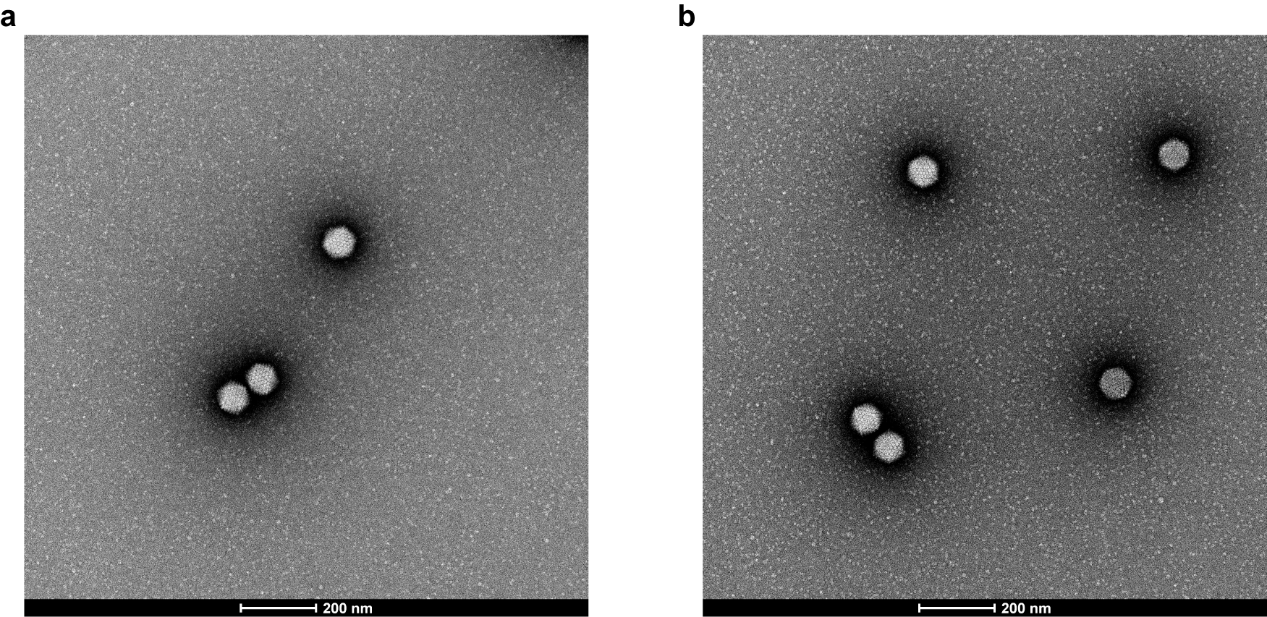

**Supplementary information, Fig. S1 Negative staining electron microscopy of Ad5-nCoV vaccines.**

Negative staining electron micrographs of **a** Ad5-nCoV_Wu vaccine particles and **b** Ad5-nCoV_O vaccine particles.

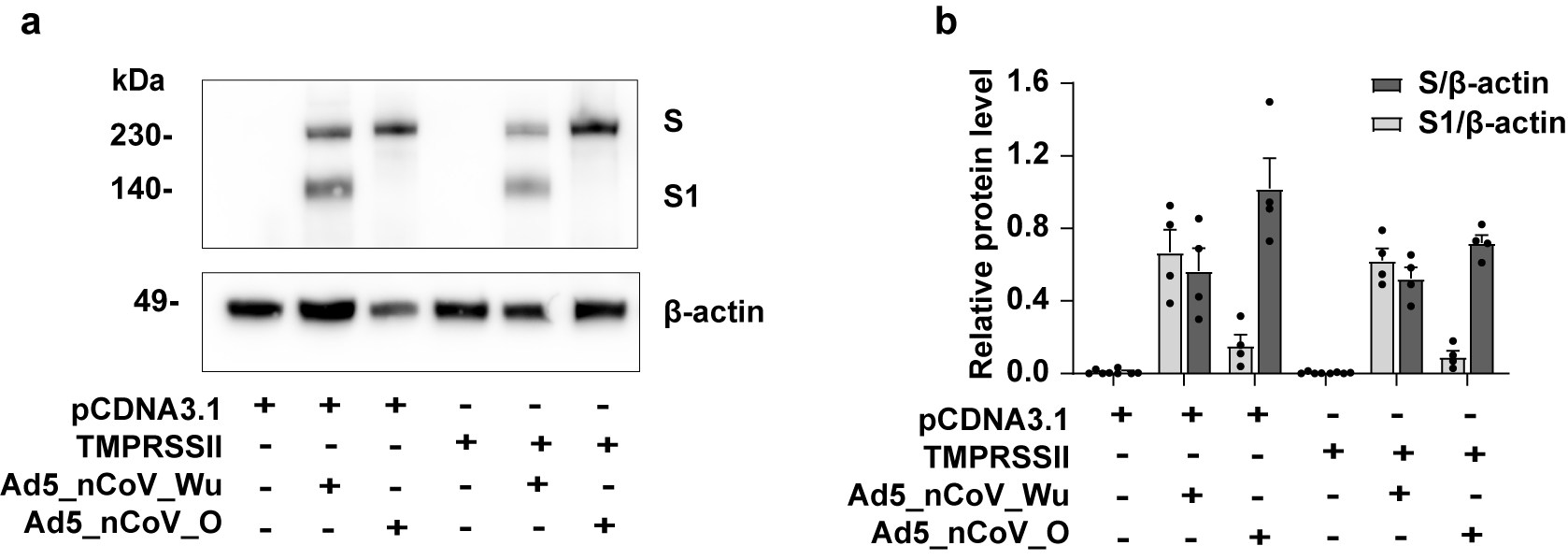

**Supplementary information, Fig. S2 Western blot showing levels of S in the vaccine-inoculated cells.**

**a** Western blot showing the expression level of S and shedded S1 subunits from Vero cells inoculated with Ad5-nCoV vaccines for 48h. **b** The relative protein levels analyzed with ImageJ. Data are represented as mean with SEM. (four biologically independent samples per group).

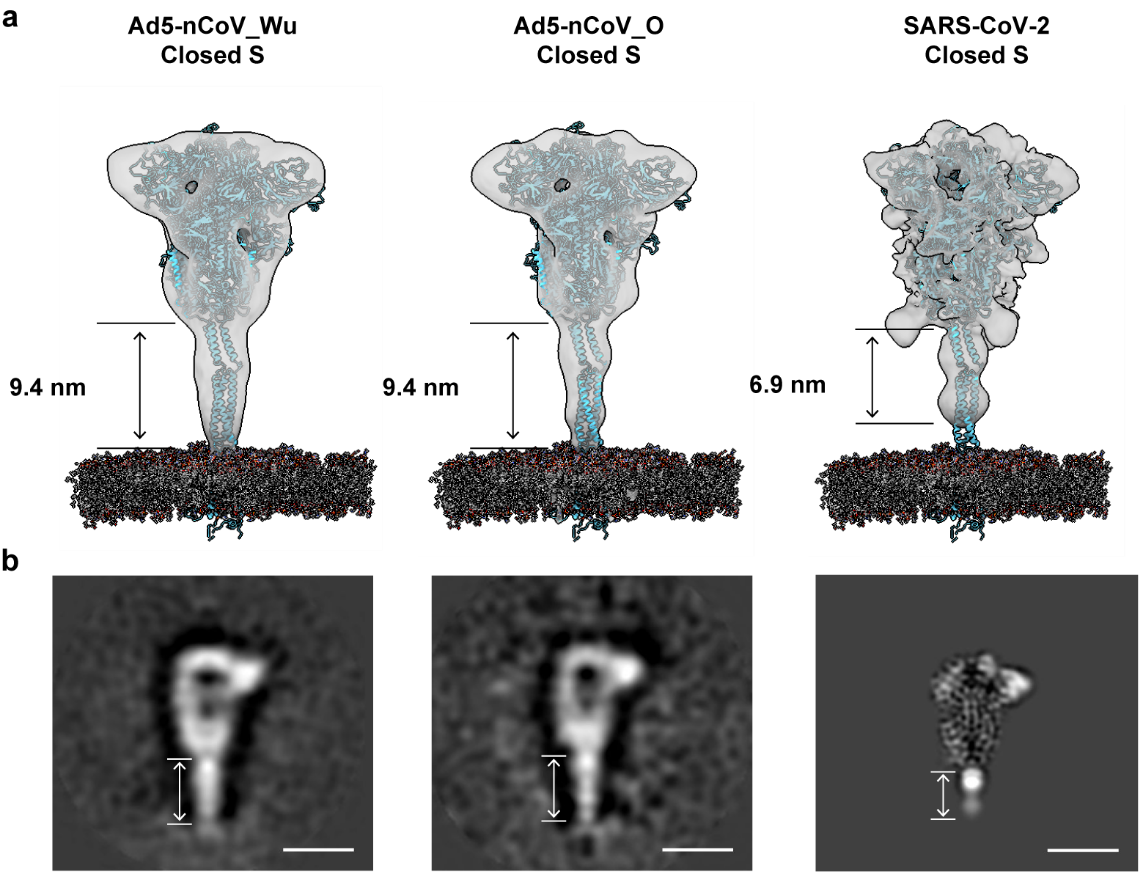

**Supplementary information, Fig. S3 The stalk region of Ad5-nCoV vaccine-induced S in closed conformation.**

**a** Maps of the closed S induced by Ad5-nCoV_Wu, Ad5-nCoV_O or from SARS-CoV-2 virion surface fitted with a full-length model of the SARS-CoV-2 closed S predicted by all-atom molecular dynamics simulation. The measured lengths of stalk regions in each map are labeled respectively. **b** The corresponding measured region in each map displayed in IMOD. Scale bar: 10 nm.

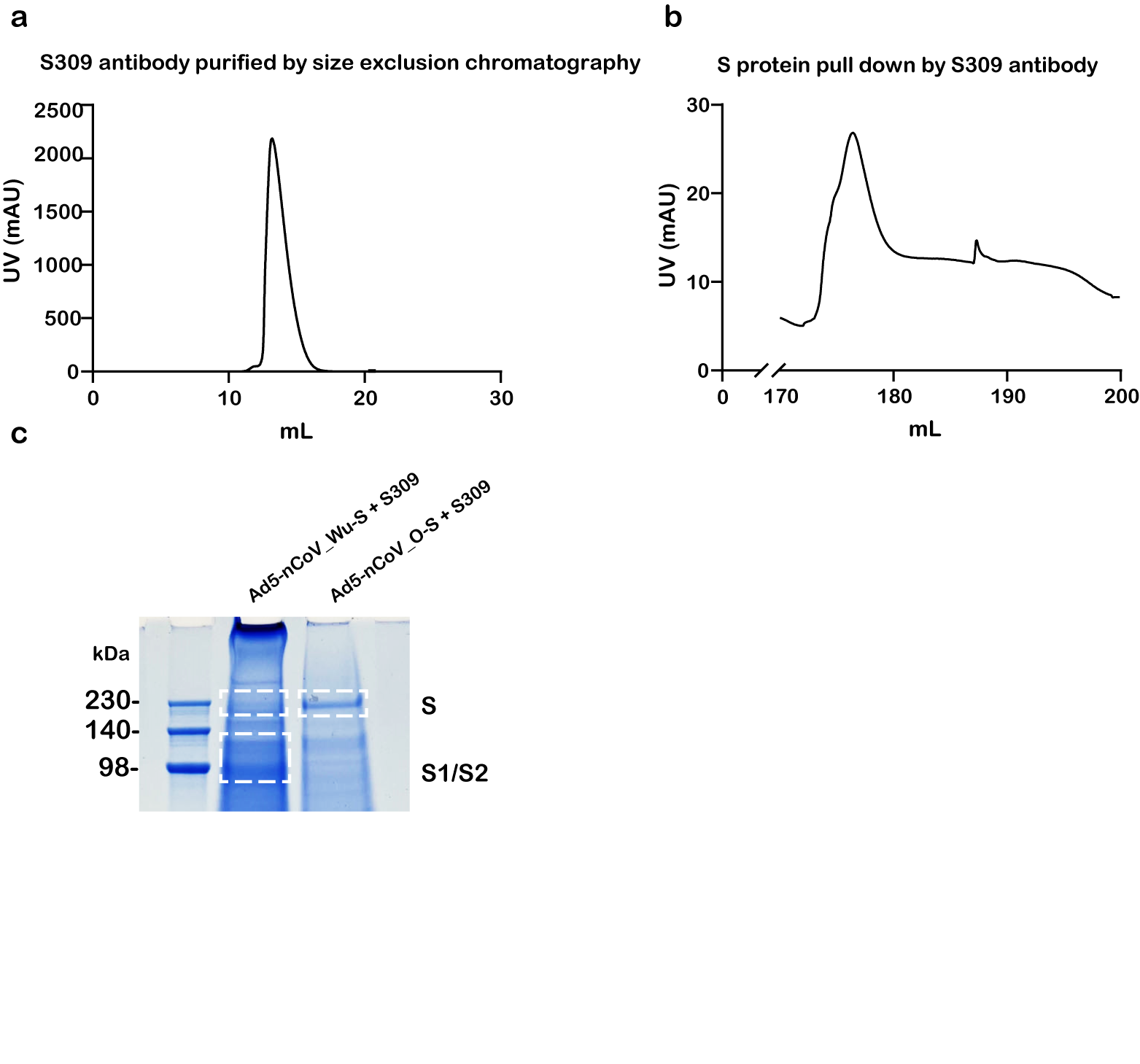

**Supplementary information, Fig. S4 Purification of S from cell membrane.**

**a** Size-exclusion chromatography of S309 antibody using Superdex 200 increase 10/300 GL column. **b** S protein pull-down assay with S309 antibody using Hitrap ProteinA HP column. **c** SDS-PAGE of spike-antibody complexes. Lane 1: protein ladder; lane 2: spike induced by Ad5-nCoV_Wu vaccine with S309 antibody; lane 3, spike induced by Ad5-nCoV_O vaccine with S309 antibody.

**Supplementary information, Table S1. Cryo-ET data collection and reconstruction statistics.**

|  | **Ad5-nCoV_Wu** | | **Ad5-nCoV_O** | |
| --- | --- | --- | --- | --- |
| **Data collection** | | | | |
| Microscope | Titan Krios | | Titan Krios | |
| Voltage (kV) | 300 | | 300 | |
| Detector | Gatan K3 | | Gatan K3 | |
| Magnification | 18,000 | | 53,000 | |
| Energy filter | NA | | Gatan BioQuantum, 20 eV | |
| Pixel size (Å) | 0.78 (super-resolution) | | 0.83 (super-resolution) | |
| Tilt schemes | Dose-symmetric scheme | | Dose-symmetric scheme | |
| Exposure (e^-^/Å^2^) | 131.2 | | 106.6 | |
| Defocus range (μm) | -4.0 ~ -5.0 | | -3.4 ~ -5.6 | |
| Software | SerialEM | | SerialEM | |
| **Reconstruction** | | | | |
| Software | Dynamo | | Dynamo | |
| No. of tomograms | 22 | | 27 | |
| Dataset | Prefusion S  (Closed) | Prefusion S  (One-RBD-up) | Prefusion S  (Closed) | Prefusion S  (One-RBD-up) |
| Final no. of particles | 894 | 842 | 500 | 1,776 |
| Symmetry imposed | C3 | C1 | C3 | C1 |
| Final Resolution (Å) | 14.0 | 22.5 | 18.4 | 17.1 |
| Final pixelsize (Å) | 3.12 | 3.12 | 3.32 | 3.32 |

**Supplementary information, Table S2. N-glycoform abundances of S protein induced by Ad5-nCoV vaccines.**

Glycans on S induced by Ad5-nCoV_Wu vaccine

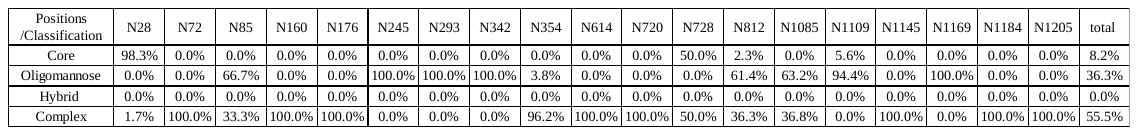

Glycans on S induced by Ad5-nCoV_O vaccine

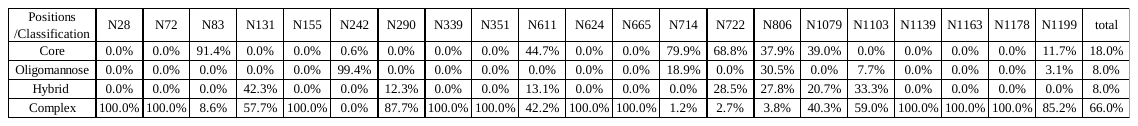

Note: The glycans on vaccines-induced S were analyzed based on their sequences starting from the signal peptide. Due to the different protein sequences and signal peptides from the WT viral S, the sequence number of each glycan site on the vaccines-induced S was slightly different from that of the WT viral S.

**Supplementary information, Table S3. Statistics of pre- and postfusion S observed on SARS-CoV-2 treated by various inactivation methods**.

| **References** | **Variants** | **Inactivation method** | **Total S/virion** | **Prefusion S/virion** | **Ratio of prefusion S** | **Postfusion S/virion** | **Ratio of postfusion S** |
| --- | --- | --- | --- | --- | --- | --- | --- |
| Yao et al., 2020^1^ | Wu | 3% PFA | 26 | 26±15 | 97% | 1±2 | 3% |
| Song et al., 2023^2^ | Delta | 3% PFA | 32 | 24±10 | 75% | 7±4 | 25% |
| Song et al., 2023^2^ | Delta | Electron-beam irradiation | 16 | 4* | 25.90% | 12±6 | 74.1% |
| Turoňová et al., 2020^3^ | D614G | 4% PFA | 40 | 39.96* | >99.9% | < 0.04* | <0.1% |
| Ke *et al.*, 2020^4^ | D614G | 4% FA | 24 | 23.28 | 97.00% | 0.72 | 3.00% |
| Liu et al., 2020^5^ | Wu | 0.05%  β-propiolactone | 7* | 1.792* | 25.6% | 5.208* | 74.4% |
| Ke et al., 2023^6^ | B.1 | 4% PFA | 21.3* | 20±10 | 93.90% | 1.3±1.5 | 6.10% |
| Ke et al., 2023^6^ | Alpha | 4% PFA | 24.4* | 23±12 | 94.26% | 1.4±1.6 | 5.74% |
| Ke et al., 2023^6^ | Gamma | 4% PFA | 28.5* | 27±13 | 94.74% | 1.5±1.8 | 5.26% |
| Ke et al., 2023^6^ | Delta | 4% PFA | 36.4* | 34±14 | 93.41% | 2.4±1.9 | 6.59% |
| Ke et al., 2023^6^ | Mu | 4% PFA | 37.8* | 35±16 | 92.59% | 2.8±2.2 | 7.41% |
| Fukuhara et al., 2023^7^ | Alpha | Active virus | 6.32* | 5.37* | 84.88% | 0.95* | 15.12% |
| Ma, X et al., 2023^8^ | BA.1 | PFA | 25.7* | 22.7±7.7 | 88.33% | 3.0±3.9 | 11.67% |
| Ma, X et al., 2023^8^ | BA.2 | PFA | 37.9* | 35.3±11.2 | 93.14 | 2.6±2.2 | 6.86% |

*Values were calculated from the reported data in references.
